# Adiponectin Preserves Metabolic Fitness During Aging

**DOI:** 10.1101/2021.01.14.426678

**Authors:** Na Li, Zhuzhen Zhang, Shangang Zhao, Yi Zhu, Christy M. Gliniak, Lavanya Vishvanath, Yu A. An, May-yun Wang, Yingfeng Deng, Qingzhang Zhu, Toshiharu Onodera, Orhan K Oz, Ruth Gordillo, Rana K. Gupta, Ming Liu, Tamas L. Horvath, Vishwa Deep Dixit, Philipp E. Scherer

**Author notes:** Children’s Nutrition Research Center, Department of Pediatric, Baylor College of Medicine, Houston, TX, USA. To whom correspondence should be addressed: Philipp E. Scherer, Ph.D., Departments of Internal Medicine, University of Texas Southwestern Medical Center, Address: L5.210, 5323 Harry Hines Boulevard, Dallas, TX 75390-8549.

## Abstract

Adiponectin is essential for the regulation of tissue substrate utilization and systemic insulin sensitivity. Clinical studies have suggested a positive association of circulating adiponectin with healthspan and lifespan. However, the direct effects of adiponectin on promoting healthspan and lifespan remain unexplored. Here, we are using an adiponectin null mouse and a transgenic adiponectin overexpression model. We directly assessed the effects of circulating adiponectin on the aging process and found that adiponectin null mice display exacerbated age-related glucose and lipid metabolism disorders. Moreover, adiponectin null mice have a significantly shortened lifespan on both chow and high-fat diet (HFD). In contrast, a transgenic mouse model with elevated circulating adiponectin levels has a dramatically improved systemic insulin sensitivity, reduced age-related tissue inflammation and fibrosis, and a prolonged healthspan and median lifespan. These results support a role of adiponectin as an essential regulator for healthspan and lifespan.

## Introduction

Healthspan and lifespan are intimately linked. Improving healthspan should help enhance the overall quality of life for an aging population, and possibly even extend lifespan (Crimmins, 2015; Piskovatska *et al*, 2019). According to current estimates, by 2050, the number of older adults in US, above the age 65 years are expected to double, rising from 40.2 million to approx. 88 million (https://www.cdc.gov/nchs/products/databriefs/db106.htm). In the U.S., the average lifespan is around 79.3 years, while the estimated healthspan is only 67.3 years, indicating that the individuals will on average live up to 20% of their lives in an unhealthy state (Olshansky, 2018). Moreover, 35-40% of adults aged 65 and above are obese. Given both aging and obesity are independent risk factors for chronic diseases, it is important to further determine how the confluence of adiposity and aging impacts healthspan and lifespan. The primary health problems associated with elderly individuals are obesity and associated metabolic disorders, including insulin resistance, type 2 diabetes, non-alcoholic fatty liver disease, hypertension, cardiovascular disease, and many types of cancers. These diseases are global public health problems, significantly accelerating the aging process, and severely decreasing the quality of life and overall life expectancy (Jura & Kozak, 2016). Thus, increasing healthspan by prolonging a disease-free period of elderly individuals may be equally important as increasing lifespan. Simple strategies, such as caloric restriction, or pharmacological interventions, such as metformin or rapamycin treatment, can promote both healthspan and lifespan in mice (Bhullar & Hubbard, 2015; Bitto *et al*, 2016; Martin-Montalvo *et al*, 2013; Minor *et al*, 2010). However, the effectiveness of such an approach in humans still awaits confirmation. The search for novel and effective strategies to extend these processes is still one of the major goals of geroscience research.

Adiponectin was one of the earliest adipokines described (Scherer *et al*, 1995). Since its discovery, significant efforts have been made to study its regulation, biogenesis, and physiological effects. As an excellent biomarker for mature adipocytes, circulating adiponectin levels are inversely correlated with fat mass, distinguishing it from most of the other adipokines, including leptin (Hu *et al*, 1996). Adiponectin exerts pleiotropic effects, including improving glucose tolerance, increasing insulin sensitivity, enhancing lipid clearance, and reducing systemic inflammation and tissue fibrosis (Scherer, 2006). Our previous studies have indicated that a lack of adiponectin in mice leads to glucose intolerance and hyperlipidemia (Nawrocki *et al*, 2006; Xia *et al*, 2018). Conversely, increasing adiponectin levels in an adiponectin transgenic mouse model, greatly improves metabolic homeostasis and produces a metabolically healthy obese phenotype (Combs *et al*, 2004; Kim *et al*, 2007). Similarly, chronic administration of adiponectin ameliorates glucose intolerance and enhances insulin sensitivity in both type 1 and 2 diabetic mice (Berg *et al*, 2001). These observations fully support the favorable effects of adiponectin in promoting metabolic health.

Most of the previous published literature focuses on beneficial effects of adiponectin in younger mice or diet-induced obese mice within less than 20 weeks of an HFD challenge. Whether similar beneficial effects could be observed in aging mice (older than 100 weeks) remains unexplored. Beyond its possible role in healthspan, some human genetics studies have implicated adiponectin as a longevity gene (Atzmon *et al*, 2008). One potential mechanism of particular interest, with robust effects on elevating circulating adiponectin levels, is the starvation hormone fibroblast growth factor-21 (FGF21). It extends lifespan in both male and female mice (Holland *et al*, 2013). Similarly, thiazolidinediones (TZDs), agonists of the peroxisome proliferator-activated receptor γ(PPARγ), also significantly increase circulating adiponectin levels, and ameliorate aged-related tissue function decline (Viljoen & Sinclair, 2009; Yu *et al*, 2002). In addition, female mice harbor higher circulating adiponectin levels and live longer compared to male mice (Gehrand *et al*, 2016). All these observations point to a positive correlation between high circulating adiponectin and longevity and implicate adiponectin as a novel circulating hormone that may directly promote both healthspan and lifespan in mice. To test this hypothesis, we used our established mouse models of adiponectin overexpression and complete absence of adiponectin and assessed the effect of circulating adiponectin on the aging process. Our results reveal that adiponectin null mice have a significantly reduced healthspan and lifespan, while adiponectin transgenic mice have a significantly prolonged healthspan.

## Methods

### Animals experiments

Adiponectin knockout mice (APN-KO) (Nawrocki *et al.*, 2006) and adiponectin transgenic mice (Combs *et al.*, 2004) with wild-type controls are on a pure C57BL6J background. All of the animal experimental protocols have been approved by the Institutional Animal Care and Use Committee of University of Texas Southwestern Medical Center at Dallas. The mice were housed under standard laboratory conditions (12 h on/off; lights on at 7:00 a.m.) and temperature-controlled environment with food and water available *ad libitum*. Mice were fed a standard chow-diet (number 5058, LabDiet, St. Louis, MO) or high-fat diet (60% energy from fat, D12492, Research Diets) for various periods as indicated in the Figures. All experiments were initiated at approximately 8 weeks of age, unless indicated otherwise. Mouse phenotyping studies were performed with controls and a minimum of two independent cohorts with more than 5 mice in each group.

### Systemic tests

Systemic tests were previously described (Zhao *et al*, 2014; Zhu *et al*, 2017). In brief, oral glucose tolerance tested were performed on overnight fasted mice. The mice orally received 1.25g or 2 g of glucose per kg body weight dissolved in phosphate buffered saline (Cat. 806552, Sigma-Aldrich). Injection volume was calculated based on 10 μl/g body weight. Blood glucose concentrations were measured by glucose meters (Contour) at the indicated time points. For ITTs, mice were fasted for 6 h in the morning, and chow-fed animals were intraperitoneally injected with insulin at a dose of 0.5 U per kg body weight, while HFD-fed animals were injected with a dose of 0.75 U per kg body weight.

Blood glucose concentrations were measured by glucose meter at the indicated time points; For T.G. clearance, mice were fasted (16 h), then gavaged 15 ul g^−1^ bodyweight of 20% intralipid (Fresenius Kabi Clyton, L.P.). Blood was collected at timed intervals then assayed for T.G. levels (Infinity; Thermo Fisher Scientific) and FFA levels (NEFA-HR); Wako Pure Chemical Industries). For some of the experiments, area under curve (AUC) was calculated.

### Blood parameters

Blood was taken from fed animals in the morning and was centrifuged at 8000 g for 5 min, and then the supernatants were collected for multiple analyses. Adiponectin was measured using an ELISA kit from Invitrogen (Catalog number: EZMADP-60K). Serum insulin levels were measured using ALPCO Mouse Insulin ELISA Jumbo kit (Cat. Number: 80-INSMS-E10). Mercodia Developing Diagnostic). Serum IGF-1 levels were measured by Mouse/Rat IGF-1 Quantikine ELISA kit (R&D Systems, Inc., Minneapolis, MN, USA). Serum parameters were measured and calculated with a VITROS analyzer (Ortho Clinical Diagnostics) at UTSW metabolic core.

### RT-qPCR and Analysis

RNA was extracted from fresh or frozen tissues by homogenization in TRIzol reagent (Invitrogen) as previously described (Zhu *et al*, 2016). We used 1 μg RNA to transcribe cDNA with a reverse transcription kit (Bio-Rad). Most of RT-qPCR primers were from the Harvard Primer Bank (https://pga.mgh.harvard.edu/primerbank/). The relative expression levels were calculated using the comparative threshold cycle method, normalized to the housekeeping gene *Gapdh*.

### Histological Analyses

For all histological analyses, four sections from at least three mice per group were stained and the examiner, typically a pathologist, was blinded to the genotype and/or treatment condition, as previously described (Zhao *et al*, 2020). In brief, for immunohistochemistry (IHC), tissues were fixed in 4% paraformaldehyde and embedded in paraffin. Sections (5 μm) were deparaffinized, heat retrieved (buffer with 10 mM Tris, 1.0 mM EDTA, PH=8.0, 94–96 °C for 30min, cool naturally), perforated (0.2% Triton × 100, 10 min), blocked in 3% BSA (Sigma, A9418) and then incubated with Mac2 (1:500 dilution, Cat#: 125401, BioLegend) primary antibodies. IHC and Hematoxylin (Vector, H3401) and Eosin Y (Thermo, 6766007) staining (HE staining) were performed using standard protocols or under the manufacturer’s instructions. Detection of IHC signal was performed with Vectastain Elite ABC kit (Vector Laboratories, Burlingame, CA) and DAB substrate kit for peroxidase (Vector Laboratories) followed by hematoxylin counterstaining (Vector Laboratories). For immunofluorescence of perilipin (1:500 dilution NB100-60554, Novus),Mac2, insulin (1:500, Dako #A0564) and glucagon (1:500, Invitrogen #18-0064), after incubation with primary antibody, slides were washed and incubated with Secondary antibodies (1:250 dilution) used were Alexa Fluor 488 or 594 donkey anti-rabbit IgG (HCL), Alexa Fluor 488 or 594 donkey anti-goat IgG (HCL) (Invitrogen) or Alexa Fluor 488 or 594 donkey anti- guinea pig IgG (HCL)at room temperature for 1 hour, then washed and sealed with Prolong Gold antifade reagent with DAPI (Life technology P36941).

### Metabolic Cage Experiments

Metabolic cage studies were conducted using a PhenoMaster System (TSE systems) at USTW Metabolic Phenotyping Core as previously described (Zhao *et al*, 2019). Mice were acclimated in temporary holding cages for 5 days before recording. Food intake, movement, and CO_2_ and O_2_ levels were measured at various intervals (determined by collectively how many cages were running concurrently) for the indicated period shown on Figures.

### Statistics

All values are expressed as the mean ± SEM. The significance between the mean values for each study was evaluated by Student t tests for comparisons of two groups. One way or two-way ANOVA was used for comparisons of more than two groups. The box-and-whisker analysis was performed to exclude potential outliner data accordingly. *P* ≤ 0.05 is regarded as statistically significant. For lifespan analysis, data were calculated using the GraphPad Prism 7 and OASIS 2 software. Log-rank (Mantel–Cox) tests were used to analyze Kaplan–Meier curves.

## Results

### Altered adiponectin levels in adiponectin null(APN-KO) and adiponectin overexpressing transgenic mice (ΔGly)

Male adiponectin null mice (APN-KO) (Nawrocki *et al.*, 2006) and adiponectin transgenic (ΔGly) mice (Combs *et al.*, 2004) were used for this study. The initial number of mice for each group in the study and a detailed scheme of the phenotypic assessments performed is outlined in (**Fig. S1A-B**). APN-KO were challenged with chow (NCD) or high-fat diet (HFD). ΔGly mice were challenged with chow diet (NCD). Consistent with expectations, serum adiponectin was absent in APN-KO mice (**Fig. S1C**). For ΔGly transgenic mice, circulating adiponectin levels were increased by 50% (**Fig. S1C**). All these observations indicate that our loss and gain of function mouse models indeed alter circulating adiponectin levels effectively as expected.

### Deletion of adiponectin in aged mice shortens lifespan on HFD

Given that the loss of adiponectin leads to impaired glucose tolerance and lipid clearance, we wanted to test whether these mice have a shortened lifespan. A cohort of APN-KO and WT mice was used to measure the lifespan. The survival curves for APN-KO reveal a statistically significant shortened lifespan compared to WT control both in the chow diet cohort (**Fig. 1A**) and in the HFD cohort (**Fig. 1B**). Thus, loss of adiponectin in mice accelerates the aging process and shortens lifespan.

**Fig. 1:**
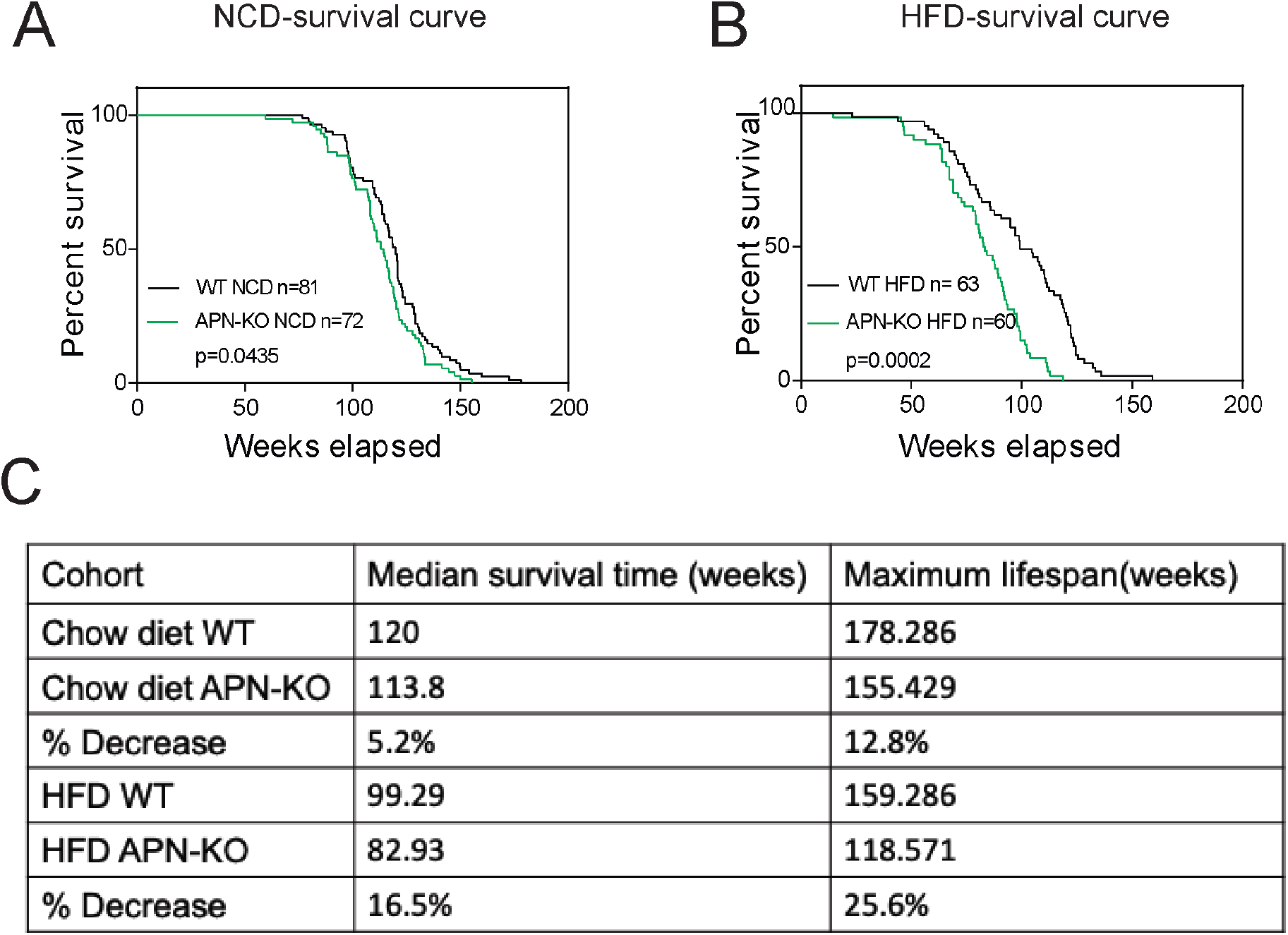
Lack of APN in aging mice shortens lifespan. A. Kaplan-Meyer survival curves for WT and APN-KO mice on chow diet. B. Kaplan-Meyer survival curves for WT and APN-KO mice on HFD. C. Median survival time and maximum lifespan for each cohort. *n* denotes the number of mice per group. *P* values were determined by log-rank (Mantel–Cox) test.

### Loss of adiponectin impairs glucose and lipid homeostasis during aging

Glucose intolerance is a hallmark of the aging process (DeFronzo, 1981). Compared to WT mice, APN-KO mice did not show any striking difference in body weight at middle- and advanced-aged, both on chow diet and on HFD (**Fig. 2A-B**). We examined glucose homeostasis in aged mice (100 weeks for with the HFD cohort and 140 weeks for the chow diet cohort). In accordance with previous metabolic studies of young adiponectin null mice, differences in glucose tolerance were marginal in mice fed standard chow diet (**Fig. 2C**). APN-KO mice fed HFD, in contrast, exhibited significantly higher glucose excursions during an OGTT (**Fig. 2D**) reflecting impaired glucose tolerance. However, no significant difference in plasma insulin level was observed during the OGTT at the different time points (**Fig. S2A**). This indicates that APN-KO mice are more susceptible to diet-induced insulin resistance.

**Fig. 2:**
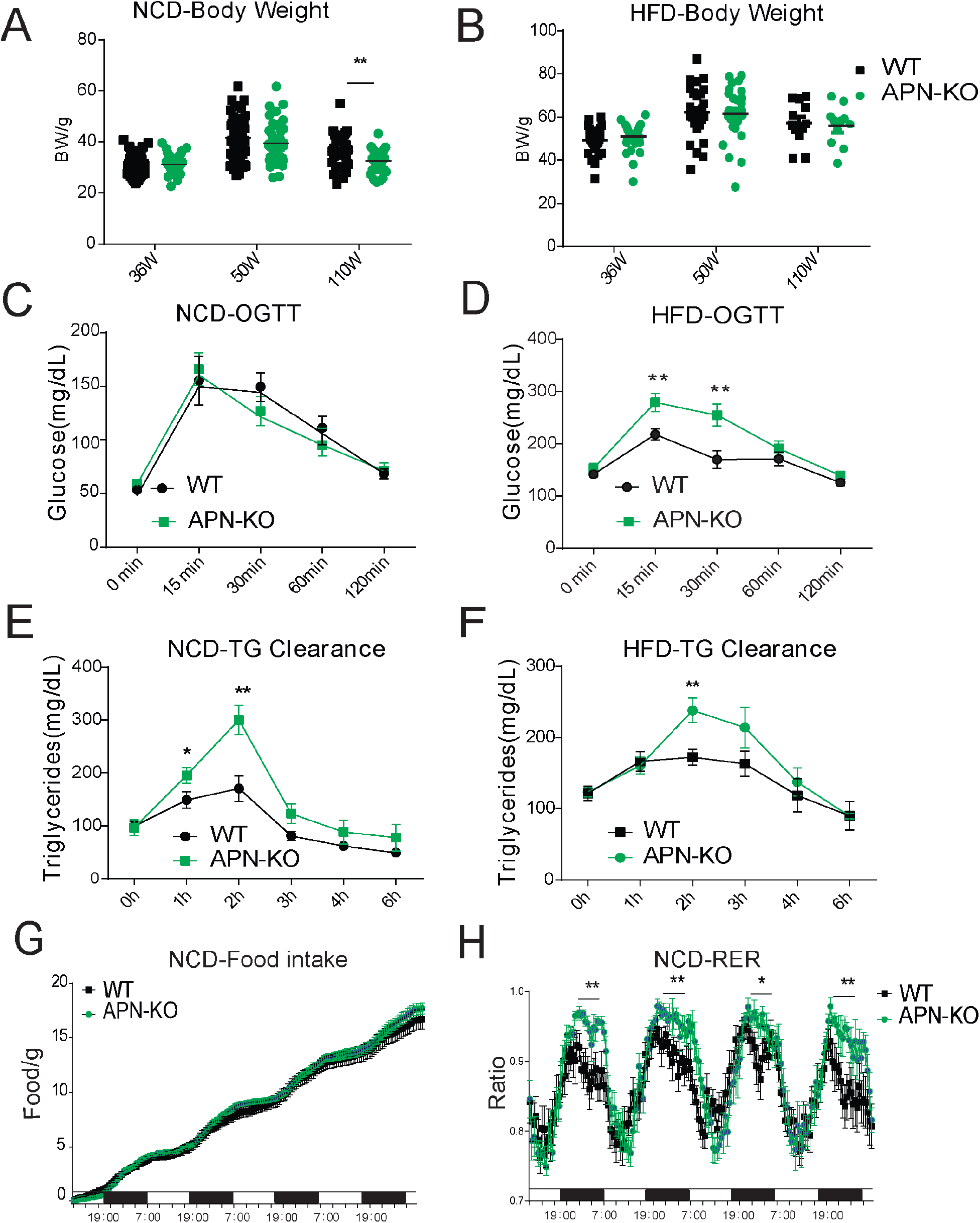
Lack of APN in aging mice attenuates glucose and lipid homeostasis. A. Body-weights during aging processes for WT and APN-KO mice fed on chow diet. B. Body-weights during aging processes for WT and APN-KO mice fed on HFD. C. An OGTT (2g kg^−1^ bodyweight; single gavage) on chow diet-feeding WT and APN-KO mice at 110-week old (n=7 per group). D. An OGTT (1.25 g kg^−1^ bodyweight; single gavage) on HFD-feeding WT and APN-KO mice at 85-week old (n=8 for WT, n=7 for APN-KO mice). E. T.G. clearance test (20% intralipid; 15 ul g^−1^ bodyweight; single gavage) in chow diet-feeding WT and APN-KO mice at 110-week old (*n* = 9 for WT, n=10 for APN-KO mice). F. T.G. clearance test (20% intralipid; 15 ul g^−1^ bodyweight; single gavage) in HFD-feeding WT and APN-KO mice at 85-week old (n=8 per group). G. Metabolic cage analyses showing food intake for chow diet-feeding WT in APN-KO mice at 110-week old (n=8 for WT, n=7 for APN-KO mice). Data are mean ± SEM. Student’s *t* test: \**p* < 0.05, \*\**p* < 0.01, \*\*\**p* < 0.001 for WT vs *APN-KO.* H. Metabolic cage analyses showing respiratory exchange rate (RER) chow diet-feeding WT and APN-KO mice at 110-week old (n=8 for WT, n=7 for APN-KO mice). Data are mean ± SEM. Student’s *t* test: \**p* < 0.05, \*\**p* < 0.01, \*\*\**p* < 0.001 for WT vs *APN-KO.*

To elucidate the effects of adiponectin on lipid metabolism of aged mice, we performed a triglyceride (TG) clearance test by gavaging the WT and APN-KO mice with 20% intralipid. Triacylglycerol levels in both NCD and HFD-fed APN-KO mice peaked at higher levels and showed a slower clearance of lipids from plasma (**Fig. 2E-F**). This highlights a prevailing impaired lipid clearance in APN-KO mice. Furthermore, although APN-KO and WT mice consume comparable amounts of diet (**Fig. 2G**), indirect calorimetry studies show that APN-KO mice had a significantly higher respiratory exchange ratio (**Fig. 2H**), indicative of carbohydrate being a more predominant fuel source in the absence of adiponectin. Combined, these results suggest adiponectin is necessary to maintain proper lipid homeostasis. Lack of adiponectin prompts glucose metabolism to be more prevalent.

### Deletion of adiponectin in aged mice exacerbates tissue functional decline

The aging process is associated with gradual decline and deterioration of functional properties at the tissue level. In aging adipose tissue, this is manifest as expansion of B cells in fat-associated lymphoid clusters(Camell *et al*, 2019), enrichment of senescent-like pro-inflammatory macrophages and loss of tissue protective macrophage subsets that drive inflammaging and compromises glucose and lipid metabolism(Camell *et al*, 2017; Lumeng *et al*, 2011; Tchkonia *et al*, 2010). In the liver and kidney, dysfunction is usually apparent as overexpression of extracellular matrix (ECM) protein constituents, such as collagen and the resulting increased fibrosis (Kim *et al*, 2016). We examined whether the deletion of APN will affect the function of these major organs. We collected adipose tissue, kidney, and liver from separate aging cohorts of young (20 weeks) and old (100 weeks for HFD cohort and 140weeks for chow diet cohort) mice. Compared to WT mice, APN-KO mice did not show significant morphological differences in adipocytes in both young and aged mice. However, the epididymal fat of APN-KO mice fed either HFD or chow diet show increased pro-inflammatory-like macrophages in the aged mice, as demonstrated by a prominent signal for the macrophage marker Mac2 (**Fig. 3A-B**). This demonstrates that the loss of adiponectin accelerates adipose tissue inflammation, a characteristic marker of the increased aging process. We do not know whether these macrophages originate from bone marrow-derived monocytes that infiltrate the tissue or whether the lack of adiponectin enhances differentiation of proliferating tissue resident monocytes into macrophages.

**Fig. 3:**
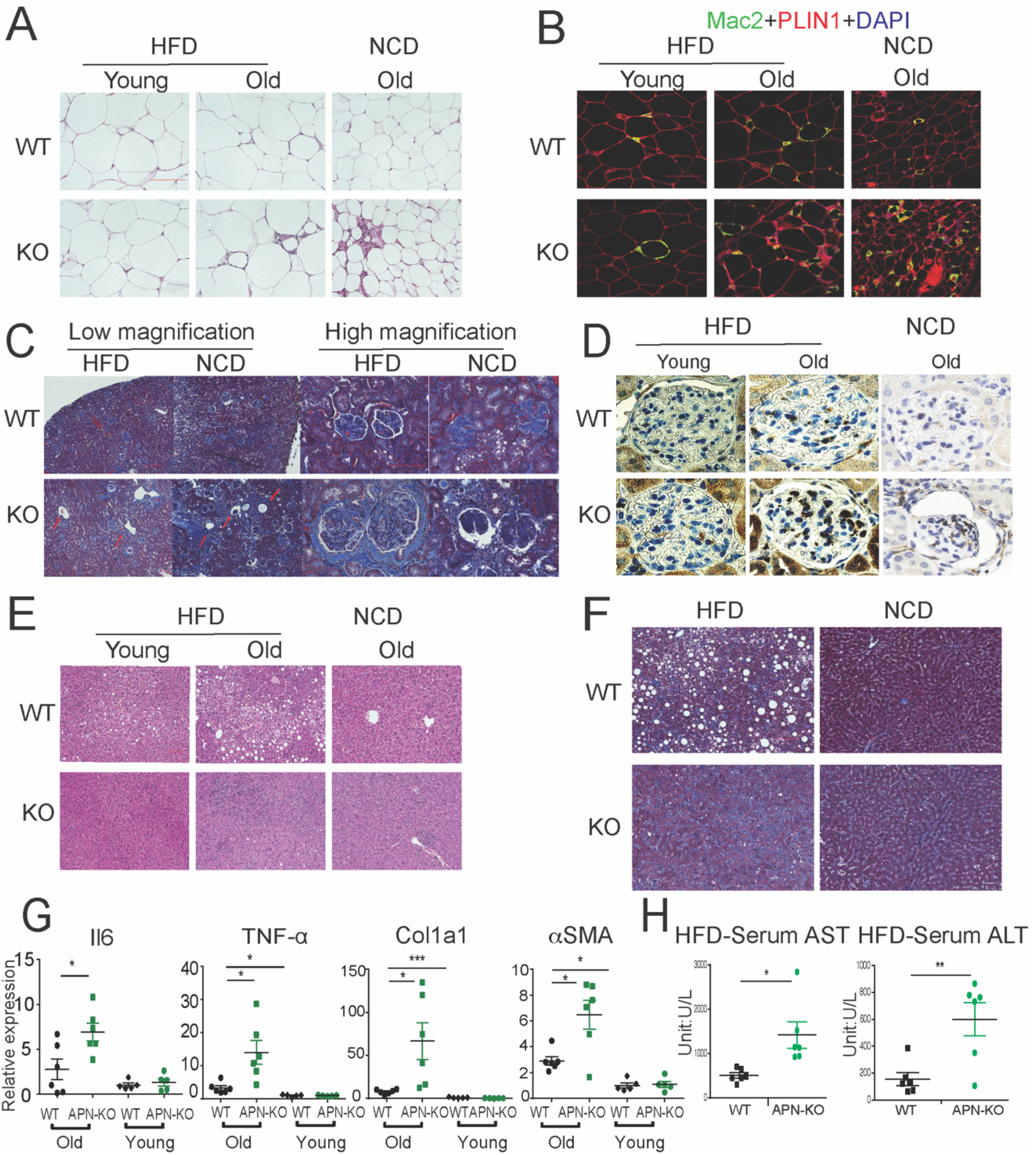
Deletion of APN in aged mice exacerbate functional decline. A. H&E staining of an Epi fat depot of 20-week old and 100-week old WT and APN-KO mice fed on HFD or 140-week old WT and APN-KO mice on chow diet. B. Mac2 staining of an Epi fat depot of 20-week old and 100-week old WT and APN-KO mice fed on HFD or 140-week old WT and APN-KO mice on chow diet. C. Trichrome staining of kidney sections reveals severe interstitial and periglomerular fibrosis in 110-week old APN-KO mice fed on HFD and 140-week old APN-KO mice fed on chow diet. Collapsed tufts are seen inside widened Bowman’s capsules forming glomerular cysts (red arrow) D. Mac2 staining of kidney sections of 20-week old and 100-week old WT and APN-KO mice fed on HFD or chow diet. E. H&E staining of Liver of 20-week old and 100-week old WT and APN-KO mice fed on HFD, 140-week old WT and APN-KO mice on chow diet. Note the extensive inflammatory cell infiltrates in the liver of the aged APN-KO mice fed on HFD. F. Trichrome stains of liver sections from 20-week old and 100-week old WT and APN-KO mice fed on HFD or 140-week old WT and APN-KO mice on chow diet, examine liver fibrosis. G. Expression of inflammatory and fibrosis markers in liver tissues of 20-week old and 100-week old WT and APN-KO mice fed on HFD or chow diet (n=5 per groups of young cohorts, n=6 per groups of aged cohorts). H. Serum AST and ALT activities in 100-week old WT and APN-KO mice fed on HFD (n=6 per group). Bar, 100μm.Data are mean ± SEM. Student’s *t* test: \**p* < 0.05, \*\**p* < 0.01, \*\*\**p* < 0.001 for WT vs *APN-KO.*

We also examined the age-related decline of health parameters in two other vital organs, kidney and liver. Even during normal aging, the kidney develops age-related structural changes and displays functional declines, including nephrosclerosis, loss of renal mass or compensatory hypertrophy of the remaining nephrons, with a corresponding decrease in glomerular filtration rate (GFR) and renal blood flow RBF (Weinstein & Anderson, 2010). Clinical studies have demonstrated that adiponectin is elevated in patients with chronic kidney disease, suggesting a possible compensatory upregulation to alleviate further renal injury (Christou & Kiortsis, 2014). Morphologically, APN-KO mice fed either the HFD or the chow diet show more severe interstitial and periglomerular fibrosis. Compared to aged WT mice, the glomeruli in aged APN-KO mice have collapsed tufts, accompanied by hypertrophic Bowman’s capsules (**Fig. 3C**). Meanwhile, aged APN-KO mice exhibited a significant increase in kidney weight as compared with aged WT mice (**Fig. S2C**). To determine the cause of this severe glomerular and tubulointerstitial damage in APN-KO mice, we investigated the glomerular infiltration with macrophages. Immunohistochemical staining with Mac2 antibodies reveals a significant increase in Mac-2 positive intraglomerular signal in the old mice which is vastly more abundant in the APN-KO mice fed the HFD (**Fig. 3D**).

Aging increases the susceptibility of various liver diseases as well, responsible for a deteriorated quality of life in the elderly and increasing mortality rate. Several studies suggest that hypoadiponectinemia predicts liver fibrosis and accelerates hepatic tumor formation (Park *et al*, 2015). Thus, we explored whether the lack of adiponectin may exacerbate age-induced dysfunction and dysmorphology of the liver. Unlike other diet-induced obese mouse models, we did not find any enhanced lipid droplet accumulation in the livers of APN-KO mice compared to WT mice upon short term and long-term exposure to HFD treatment (**Fig. 3E**). However, we found many inflammatory infiltrates in the livers of APN-KO mice on HFD diet. The expression of inflammatory markers is significantly increased in aged APN-KO mice fed on HFD and chow diet (**Fig. 3G, Fig. S2B**), indicative of increased inflammation in the liver. Moreover, Trichrome staining highlighting the ECM reveals increased hepatic fibrosis in old APN-KO mice on the chow diet that was even more evident under HFD conditions (**Fig. 3F**). Mirroring these histological findings, the expression levels of liver fibrosis markers, such as Col1α1 and αSMA, are strikingly increased in older HFD and chow diet fed APN-KO mice (**Fig. 3G, Fig. S2B**). Liver damage was further confirmed by elevated serum AST and ALT levels in HFD fed APN-KO mice compared with control mice (**Fig. 3H**). All of these observations support that adiponectin plays an essential role in maintaining normal liver function during the aging process.

Upon comparing young WT vs APN-KO mice (20 weeks) that were exposed for 8 weeks to HFD, no genotype-specific differences were observed in the kidney and the liver. This therefore indicates that the pathological changes in older APN-KO mice genuinely reflect age-related chronic changes rather than simple developmental differences that would be apparent in the young mice as well. These findings clearly indicate that the lack of adiponectin during aging exacerbates liver and renal damage, at least in part through proinflammatory mechanisms.

### Increasing adiponectin protects mice from aged induced metabolic dysfunction

Clinically, adiponectin levels are significantly higher in centenarians and in some of their offspring, suggesting that adiponectin may be a key driver to promote healthspan and lifespan. As the elimination of adiponectin shortens healthspan and lifespan, we wondered whether increasing adiponectin by our previously established transgenic mouse model (that we refer to as the “ΔGly mouse”) could promote both healthspan and lifespan. A large cohort of WT and ΔGly mice were placed on chow diet to assess their lifespan. After calculation, a median lifespan in Control mice was around 117 weeks, while this value in ΔGly mouse has been extended to 128 weeks (9% extension), indicating that increasing circulating adiponectin prolongs median lifespan. However, the maximum lifespan is comparable in Control and ΔGly mice, as the overall survival curves were not different by log rank test (**Fig. 4A**).

**Fig. 4:**
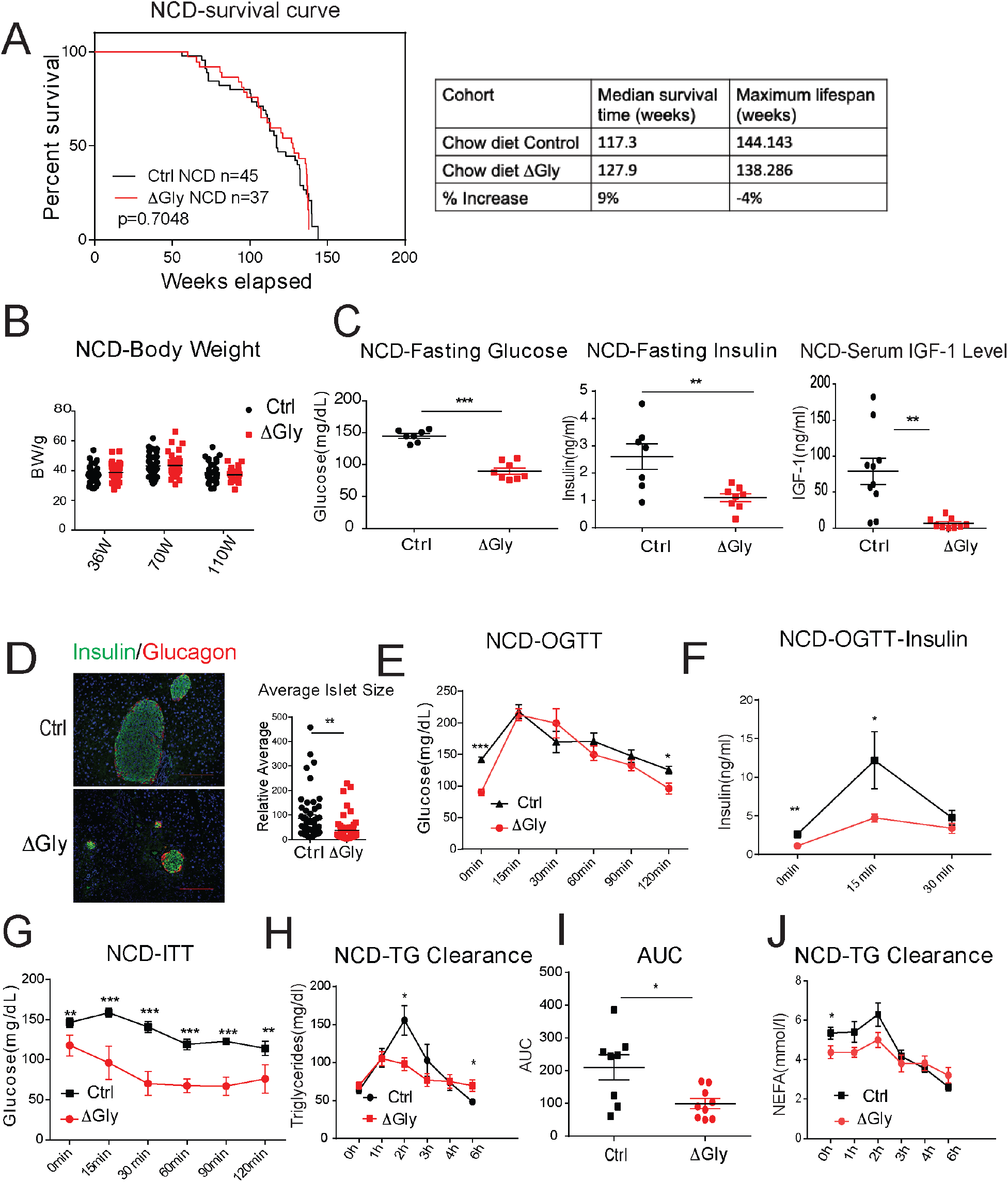
Increasing adiponectin protect against aging-induced metabolic disturbance. A. Kaplan-Meyer survival curves for WT and ΔGly mice on chow diet. Median survival time and maximum lifespan for each cohort. *n* denotes the number of mice per group. *P* values were determined by log-rank (Mantel–Cox) test. B. Body-weights during aging processes for controls and ΔGly mice fed on chow diet. C. Systemic glucose, insulin and IGF-1 levels in 50-week old controls and ΔGly mice after fasting 16h. D. Insulin and glucagon IF staining of pancreases from controls and ΔGly mice at 140-week old (left). Right: Relative average islet size. E. An OGTT (2 g kg^−1^ bodyweight; single gavage) revealed marginally improved glucose tolerance in 50-week ΔGly compared with controls (n=8 per group). F. Serum insulin levels during glucose tolerance test performed in panel C (n=8 per group). G. ITT in controls and ΔGly mice at 50-week old. (n=8 per group) H. T.G. clearance test in controls and ΔGly mice at 50-week old (n=8 for WT, n=9 for ΔGly mice) I. AUC calculated based on H. J. Circulating FFA levels in controls and ΔGly mice at 50-week old during T.G. clearance performed in panel I. (n=8 for WT, n=9 for ΔGly mice).Bar,100μm.Data are mean ± SEM. Student’s *t* test: \**p* < 0.05, \*\**p* < 0.01, \*\*\**p* < 0.001 for controls vs ΔGly.

Besides its positive effects in prolonging median lifespan, we determined if increasing adiponectin levels may have beneficial effects in extending healthspan. Previous studies indicated that increasing adiponectin levels results in improved glucose and lipid profiles in younger mice (Berg *et al.*, 2001). However, whether these beneficial effects of adiponectin carry to older age has not been assessed. When fed with a chow diet, ΔGly mice show a similar body weight during lifespan, compared to littermate controls (**Fig. 4B**). Then we measured fasting glycemia, insulin, and IGF-1(**Fig. 4C**). Under 16hr fasted conditions, ΔGly mice have a significantly lower fasting glycemia, accompanied by a robust reduction in plasma insulin. Moreover, a reduction in circulating IGF-1 levels is observed in ΔGly mice. Lower IGF-1 levels are thought to play a key role as a mediator of health- and lifespan extension (Bartke *et al*, 2003). To test whether the improvements in systemic insulin sensitivity are also associated with improvements at the level of the pancreatic β cell, we performed H&E staining on pancreatic sections. Consistent with the reduced demand on islets to produce and release insulin in a more insulin-sensitive environment, the average islet size was considerably reduced by adiponectin overexpression, with islet structural integrity fully preserved (**Fig. 4D**). Immunohistochemical analysis of islets exhibits a normal composition with α cells (glucagon) and β cells (insulin) in ΔGly mice. During an oral glucose tolerance test, ΔGly mice displayed a much lower glucose excursion than littermates (**Fig. 4E**). In addition, insulin levels in ΔGly mice were significantly lower in response to the glucose challenge, which further supports improved insulin sensitivity (**Fig. 4F**). To confirm this, we performed insulin tolerance tests. ΔGly mice show a significant increase in insulin sensitivity (**Fig. 4G**), which is consistent with our results for the young mice. Moreover, when orally challenged with triglycerides, ΔGly mice display enhanced lipid clearance (**Fig. 4H-I**), with correspondingly lower FFA values (**Fig. 4J**). These data demonstrate that increasing adiponectin levels significantly promotes metabolic fitness in aged mice.

### Increasing adiponectin levels improves the age-related functional decline in tissues of aged mice

To probe tissue functional declines that might contribute to metabolic syndrome in the elderly, we evaluated the function of fat and liver in aged mice. Aging is associated with a redistribution of fat from the periphery to central fat deposition(Kuk *et al*, 2009). The redistribution and ectopic fat deposition with aging appear to accelerate onset of multiple age-related diseases. A histological examination of adipose tissue showed that ΔGly mice harbor much smaller adipocytes in subcutaneous and gonadal fat (**Fig. 5A**) compared to controls at the age of 140 weeks. In agreement with the epididymal adipocyte size and fat mass, inflammation is potently suppressed in visceral fat tissues of ΔGly mice, as demonstrated by a significantly reduced Mac-2 staining (**Fig. 5B**). Moreover, it was quite apparent that visceral fat pad weight was reduced in ΔGly mice with a slightly increase in subcutaneous adipose tissue (**Fig. 5C**). Aged WT mice revealed an unclear boundary in the hepatic lobule with lose cellular cytoplasm, while ΔGly mice entirely prevented lipid droplet accumulation and age-related deterioration of the morphology of the liver (**Fig. 5D**). Furthermore, gene expression of inflammation and fibrosis markers in the livers were dramatically reduced in ΔGly mice compared with their littermates (**Fig. 5E**). Combined, these findings strongly support that adiponectin promotes metabolic fitness, by maintaining a proper fat distribution, and reducing adipose tissue inflammation, along with reducing inflammation and fibrosis in liver.

**Fig. 5:**
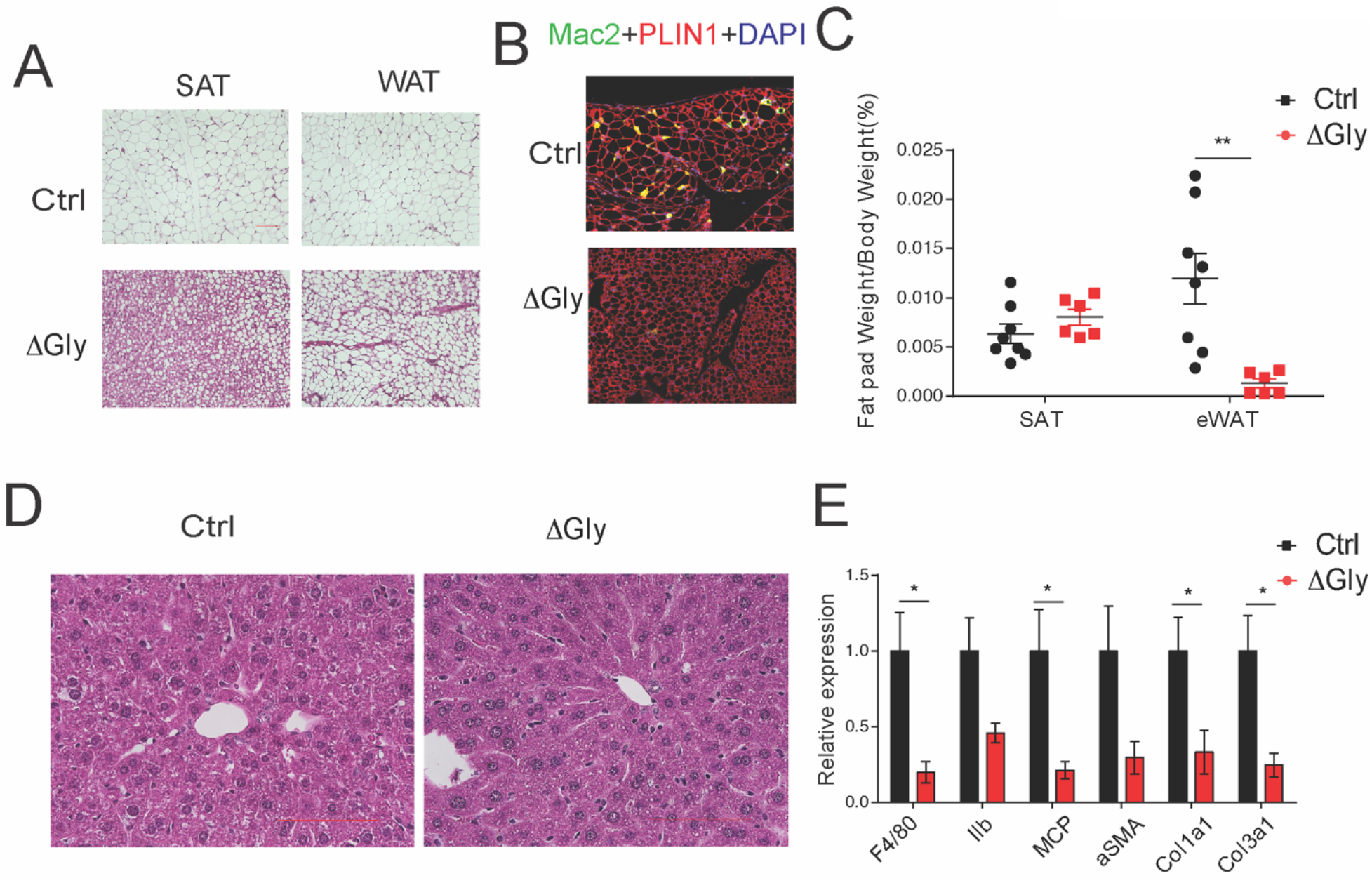
Old adiponectin overexpressing mice exhibit improved glucose and lipid homeostasis. A. H&E staining of SubQ fat depot and Epi fat depot of 140-week old controls and ΔGly mice fed on chow diet. B. Mac2 staining of Epididymal fat sections in 140-week old controls and ΔGly mice. C. Relative subcutaneous and visceral fat pad weights of 140-week old controls and ΔGly mice fed on chow diet (n=8 for controls, n=6 for ΔGly mice). D. H&E staining of Liver from 140-week old controls and ΔGly mice fed on chow diet. E. Expression of inflammatory and fibrosis markers in liver of 140-week old controls and ΔGly mice fed on chow diet (n=8 for controls, n=6 for ΔGly mice). Bar, 100μm.Data are mean ± SEM. Student’s *t* test: \**p* < 0.05, \*\**p* < 0.01, \*\*\**p* < 0.001 for WT vs ΔGly.

## Discussion

Based on data from clinical correlations as well as ample preclinical results, we appreciate that elevated levels of adiponectin are generally associated with an improved overall metabolic phenotype. Here, we systematically assessed the impact of adiponectin in the context of aging. Using adiponectin-null and adiponectin overexpressing mouse models, we have made the following observations:1) The lack of adiponectin in mice curtails healthspan by impairing glucose and lipid homeostasis, and accelerating fibrogenesis in multiple tissues, resulting in reduced healthspan; 2) The lack of adiponectin in mice shortens lifespan both on chow and HFD. 3) Increasing adiponectin levels in aged adiponectin overexpressing mice produces a healthy metabolic phenotype, with greatly increased glucose tolerance and insulin sensitivity, enhanced lipid clearance, lowered visceral fat and potent protection from inflammation and fibrosis; 4) Adiponectin overexpressing mice on a chow diet show a 9% increase in median lifespan. All these observations support that adiponectin is a vastly underestimated player in healthspan and lifespan.

With the extension of life expectancy, larger segments of the elderly population suffer from various chronic diseases. The normal aging process is associated with chronic inflammation and thereby increases susceptibility to these chronic morbidities (Goldberg & Dixit, 2015). Indeed, we observed an exacerbated pro-inflammatory state in aged WT mice compared with younger WT animals. In order to combat these age-related inflammatory changes, we need effective anti-inflammatory interventions. But these interventions should not negatively impact desired innate and adaptive immune responses. Circulating adiponectin levels are negatively correlated with inflammatory markers in diabetic patients (Krakoff *et al*, 2003; Mantzoros *et al*, 2005) and in non-diabetic subjects (Choi *et al*, 2007). Among different adipokines, adiponectin is recognized as a major adipokine regulating inflammation in a number of cell types. Our previous studies indicated acute APN depletion leads to an upregulation of inflammatory genes (Xia *et al.*, 2018). Adipose tissues are also susceptible to fibrosis. Chronic inflammation frequently results in fibrosis, which leads to functional declines in tissues. During normal aging, fibrosis occurs in small steps. Due to gradual deposition of collagens, organs become rigid and dysfunctional. Eventually, this causes the health to decline. We found indeed that aged WT mice develop more severe fibrosis compared to young WT mice.

Adiponectin has potent anti-fibrotic effects in the liver by activating peroxisome proliferator-activated receptor-gamma pathways (Shafiei *et al*, 2011), which in turn diminishes the expression of pro-fibrotic genes. Despite having reduced levels of triglyceride accumulation in the liver, the chronic lack of adiponectin dramatically exacerbates age-related liver fibrosis in parallel with disruption of liver function. In contrast, ΔGly mice are completely protected from diet- and aging- induced steatohepatitis and fibrosis, indicating of a crucial role of adiponectin in regulating liver inflammatory reactions and fibrosis. Hence, the anti-inflammatory impact and potent anti-fibrotic actions of adiponectin make it a potent novel regulator enhancing health span.

Obesity-associated chronic inflammation and insulin resistance are regarded as a pivotal risk factors for the development of several age-related pathological sequelae (Huffman & Barzilai, 2009; Kanneganti & Dixit, 2012).Improved metabolic homeostasis is positively associated with lifespan in humans and mice. Prolongevity intervention, caloric restriction and long-lived Ames dwarf mice have increased adiponectin expression (Hill *et al*, 2016). We found adiponectin mimics to some extent the impact of caloric restriction on reduction in inflammation and improved metabolic homeostasis. Reduction in adiposity is considered to be a hallmark of caloric restriction, which is an important component of its beneficial effects on metabolism. After long-term caloric restriction, aged mice display lower adiposity, smaller adipocytes and improved insulin sensitivity (Miller *et al*, 2017). Strikingly, increases in adiponectin expression are detected in these smaller adipocytes after caloric restriction. Adipocyte hypertrophy is associated with cellular stress and obesity-associated metabolic complications. Due to the limited capacity to expand subcutaneous adipose tissue in aged populations, adipocyte hypertrophy also occurs in visceral fat, which is associated with lipid spillover in multiple tissues in aging and ectopic fat accumulation(Tchkonia *et al*, 2013). Hypertrophic adipocytes and impaired redistribution of lipids exert a negative impact on insulin responsiveness, contributing to many metabolic diseases frequently observed in the elderly. Thus, metabolic disorders frequently go hand in hand with aging. Clinical studies have identified an strong inverse relationship between circulating adiponectin and insulin resistance in obese individuals (Turer *et al*, 2011). Our previous data suggested that adiponectin strongly suppresses hepatic gluconeogenesis and enhances fatty acid oxidation, thereby strongly contributing to an overall beneficial metabolic regulation (Wang & Scherer, 2016). Our aged adiponectin transgenic mice still have dramatically improved insulin sensitivity in parallel with reduced plasma insulin. All of this happens primarily due to an increase in adipocyte numbers in subcutaneous fat of ΔGly mice. In the absence of the protective effects of adiponectin, aged APN-KO mice exacerbated diet-and aging-induced glucose intolerance and lipid disorders. Moreover, ΔGly mice show improved insulin sensitivity in parallel with lower insulin and IGF-1 levels, and higher IGF-1 in APN-KO mice. Attenuated activation of the growth hormone -insulin-like growth factor I (IGF-I) axis is also an integral component of the beneficial effects of caloric restriction leading to prolonged healthspan and lifespan in rodents. Interestingly, elevated adiponectin is detected in all long-live mice: This includes the adipocyte-specific insulin receptor knockout mice (FIRKO), the Ames dwarfs (df/df) and GHRKO mice (Blüher *et al*, 2002; Masternak *et al*, 2012; Wang *et al*, 2006). These similarities across all these models suggest that an increase in adiponectin levels may be the common denominator driving longevity in all these models.

Combined, our studies and along with previous reports, demonstrate that adipose tissue plays a vital role in the aging process. In aging, dysfunctional fat tissue leads to ectopic fat deposition, lipodystrophic adipocytes, and subcutaneous fat loss, thereby contributing to increased systemic inflammation, metabolic disturbances, and functional declines in other organs. However, healthy fat pads have characteristic features not only in terms of the quantity, but more importantly, by the quality of adipose tissue (Kusminski *et al*, 2012). Adiponectin is a key player maintaining glucose and lipid homeostasis on the basis of its lipid storing capacity and its ability to communicate with other organs. Thus, overexpressing adiponectin results in the healthy expansion of subcutaneous adipose tissue, a reduction of visceral fat and improvement of inflammation and fibrosis in the liver, all of which greatly alleviates metabolic disturbances and protects against tissue functional decline during the aging process. In contrast, adiponectin deficiency increases susceptibility to metabolic diseases in the elderly. Impaired glucose tolerance and lipid clearance, severe inflammation accompanied by dysfunctional liver and kidney all reduce the quality of life and lifespan in the elderly. Thus, the ability to prolong health span by maintaining adiponectin levels provides a promising therapy for aged- related disorders and improving quality of life in older individuals.

### Study approval

The Institutional Animal Care and Use Committee of the University of Texas Southwestern Medical Center approved all animal experiments (APN:2015-101207G).

## Author contributions

NL performed most of the experiments. ZZ, SZ, YAA, YZ, LV and MW conducted some of the experiments. ZZ, YD and TO helped with the breeding of mouse models. RG performed the metabolite measurement experiment. QZ, RKG, ML and VDD gave useful suggestions to this study. NL and PES wrote the manuscript. SZ, YZ, CG and VDD revised the manuscript. PES, TLH and VDD were involved in experimental design, experiments, data analysis, and the interpretation of data.

## Acknowledgements

We thank all members of Scherer for their support of this study. We would also like to thank the UTSW Metabolic Core Facility, the Histo-Pathology Core, UTSW ARC, and Charlotte Lee for help with histology.

## Funding

This study was supported by US National Institutes of Health grant P01-AG051459 to T.L.H., V.D.D. and P.E.S.

## Competing interests

None of the authors declare any conflicts of interest.

## Supplemental Figure Legends

**Fig. S1:**
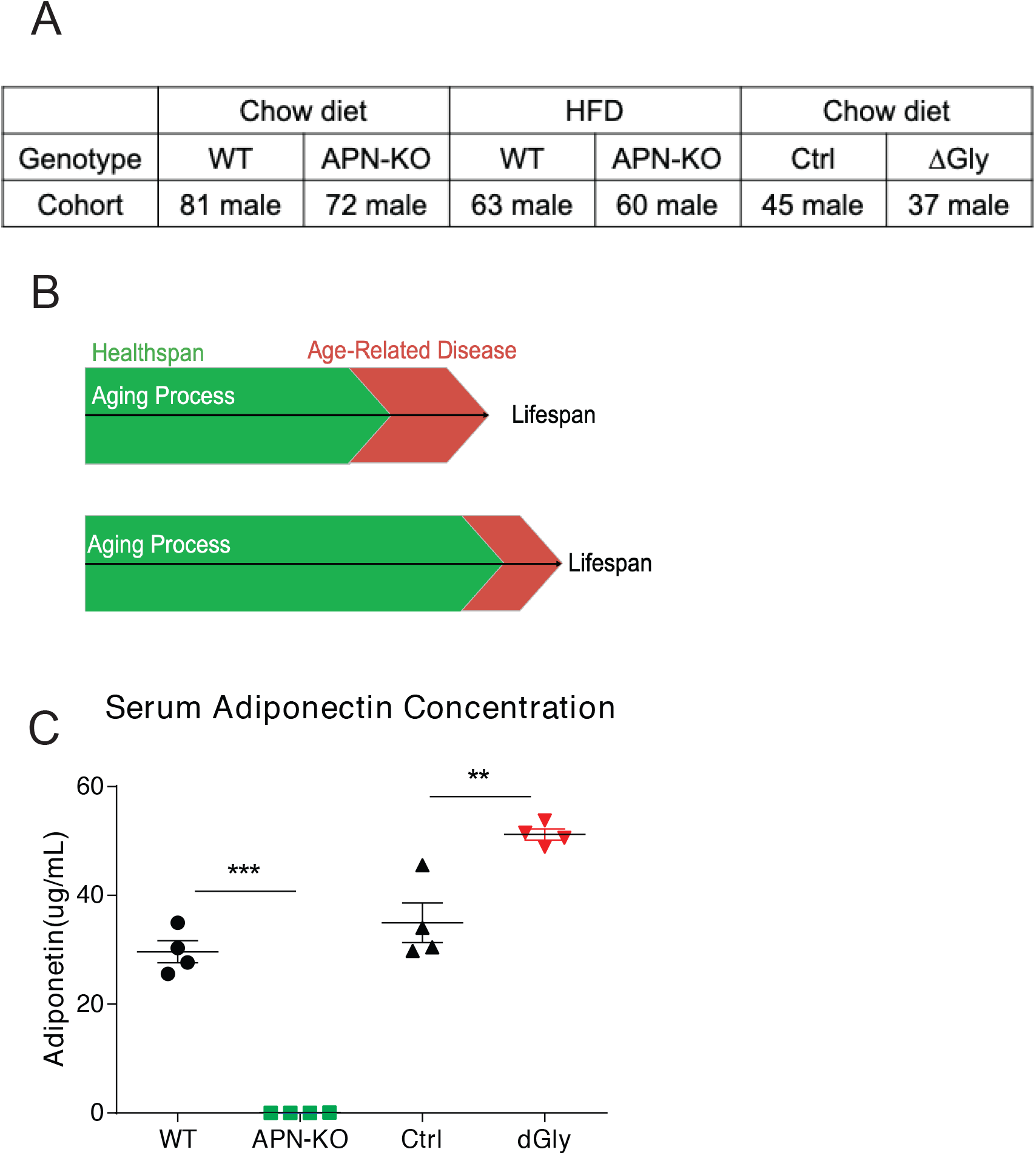
Mouse Models used for longevity studies: APN-KO Mice and ΔGly Mice. S1A. Experimental strategy for longevity experiments. S1B. Diagram of the aging process. Lifespan and healthspan are always strongly coupled. S1C. Circulating adiponectin levels measured in 50-week old APN-KO and ΔGly mice with their controls fed on chow diet respectively (n=4 per group).

**Fig. S2:**
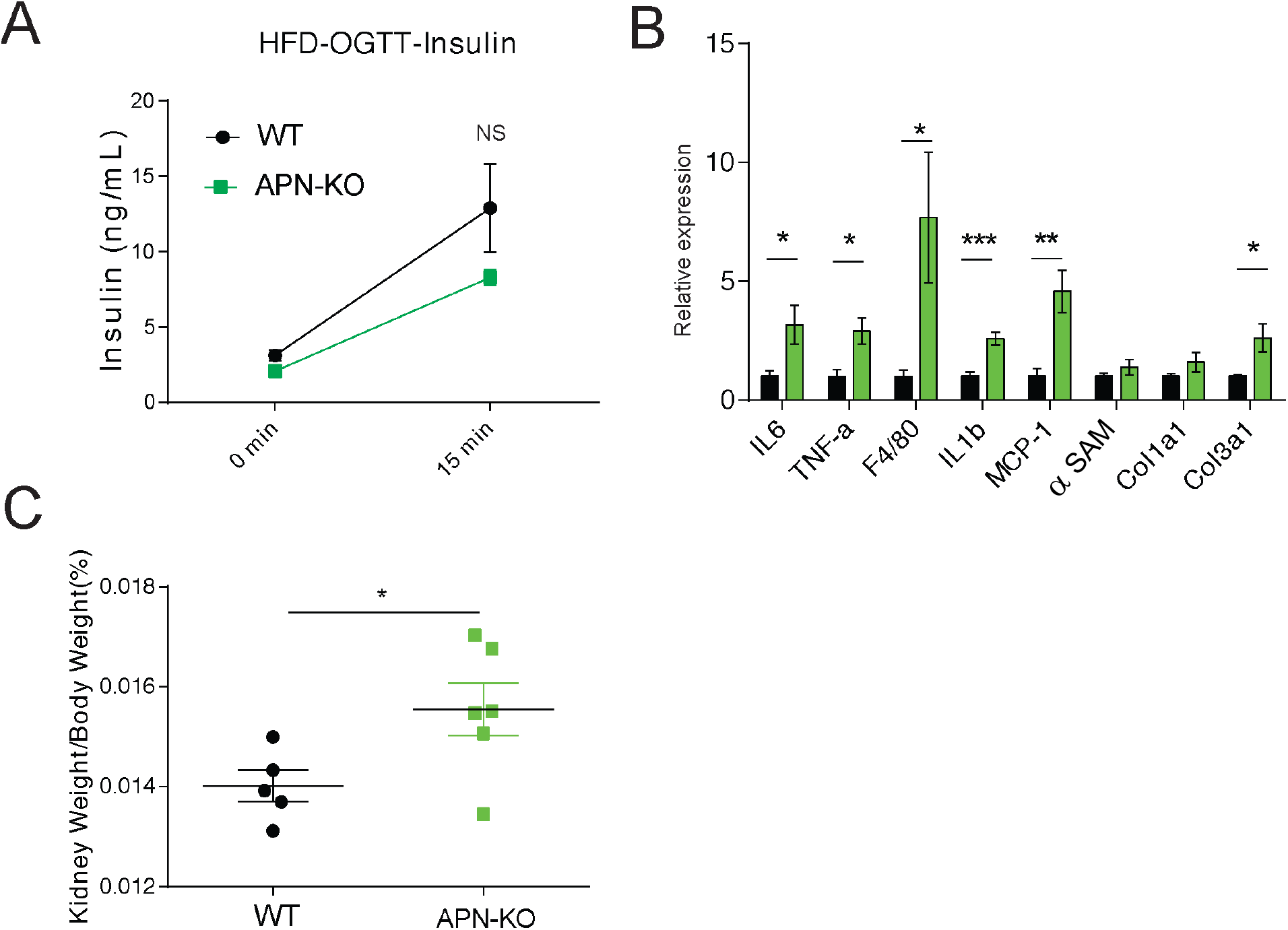
Insulin levels in APN-KO mice during OGTTs. S2A. No difference in insulin levels during OGTTs in aged APN-KO mice on HFD. And chow diet fed aged APN-KO mice do not improve glucose tolerance. Serum insulin levels during glucose tolerance test performed in Fig.3D. (n=8 for WT, n=7 for APN-KO mice. S2B. Expression of inflammatory and fibrosis markers in liver of 140-week old WT and APN-KO mice fed on chow diet (n=7 for WT, n=7 for APN-KO mice). S2C. The relative wet kidney weight with respect to body weight of 140-week old WT and APN-KO mice fed on chow diet (n=5 for WT, n=6 for APN-KO mice). Bar, 100μm.Data are mean ± SEM. Student’s *t* test: \**p* < 0.05, \*\**p* < 0.01, \*\*\**p* < 0.001 for WT vs APN-KO.

